# Dynamic competition for hexon binding between core protein VII and lytic protein VI promotes adenovirus maturation and entry

**DOI:** 10.1101/857581

**Authors:** Mercedes Hernando-Pérez, Natalia Martín-González, Marta Pérez-Illana, Maarit Suomalainen, Philomena Ostapchuk, Gabriela. N. Condezo, M. Menéndez, Urs F. Greber, Patrick Hearing, Pedro J. de Pablo, Carmen San Martín

**Affiliations:** Department of Macromolecular Structures, Centro Nacional de Biotecnología (CNB-CSIC), 28049 Madrid, Spain; Department of Condensed Matter Physics, Universidad Autónoma de Madrid, 28049 Madrid, Spain; Department of Molecular Life Sciences. University of Zurich, Winterthurerstrasse 190, CH-8057, Zurich, Switzerland; Department of Microbiology and Immunology, Renaissance School of Medicine, Stony Brook University, Stony Brook, NY, 11794-5222, USA; Instituto de Química Física Rocasolano (IQFR-CSIC), 28006 Madrid, Spain, and Ciber of Respiratory Diseases (CIBERES), ISCIII, 28029 Madrid, Spain; Instituto de Física de la Materia Condensada (IFIMAC), Universidad Autónoma de Madrid, 28049, Madrid, Spain

**Keywords:** Adenovirus, DNA virus, virus assembly, virus entry, virus maturation

## Abstract

Adenovirus minor coat protein VI contains a membrane-disrupting peptide which is inactive when VI is bound to hexon trimers. Protein VI must be released during entry to ensure endosome escape. Hexon:VI stoichiometry has been uncertain, and only fragments of VI have been identified in the virion structure. Recent findings suggest an unexpected relationship between VI and the major core protein, VII. According to the high resolution structure of the mature virion, VI and VII may compete for the same binding site in hexon; and non-infectious human adenovirus type 5 particles assembled in the absence of VII (Ad5-VII-) are deficient in proteolytic maturation of protein VI and endosome escape. Here we show that Ad5-VII- particles are trapped in the endosome because they fail to increase VI exposure during entry. This failure was not due to increased particle stability, because capsid disruption happened at lower thermal or mechanical stress in Ad5-VII- compared to wildtype (Ad5-wt) particles. Cryo-EM difference maps indicated that VII can occupy the same binding pocket as VI in all hexon monomers, strongly arguing for binding competition. In the Ad5-VII- map, density corresponding to the immature amino-terminal region of VI indicates that in the absence of VII the lytic peptide is trapped inside the hexon cavity, and clarifies the hexon:VI stoichiometry conundrum. We propose a model where dynamic competition between proteins VI and VII for hexon binding facilitates the complete maturation of VI, and is responsible for releasing the lytic protein from the hexon cavity during entry and stepwise uncoating.

**Significance Statement:** Correct assembly of an adenovirus infectious particle involves the highly regulated interaction of more than ten different proteins as well as the viral genome. Here we examine the interplay between two of these proteins: the major core protein VII, involved in genome condensation, and the multifunctional minor coat protein VI. Protein VI binds to the inner surface of adenovirus hexons (trimers of the major coat protein) and contains a lytic peptide which must be released during entry to ensure endosome rupture. We present data supporting a dynamic competition model between proteins VI and VII for hexon binding during assembly. This competition facilitates the release of the lytic peptide from the hexon cavity and ensures virus escape from the early endosome.

## Introduction

Adenoviruses are complex non-enveloped, double-stranded DNA (dsDNA) viruses. The virion of human adenovirus type 5 (HAdV-C5) consists of a 95 nm diameter icosahedral *pseudo*T = 25 protein shell enclosing a non-icosahedral DNA-protein core. The capsid is composed of 240 hexon trimers, 12 pentameric penton bases associated with a fiber sitting at each vertex, and four minor coat proteins: polypeptide IX (240 copies) on the outer capsid surface, and proteins IIIa (60 copies), VI (~320 or 360 copies, depending on two different stoichiometry determinations), and VIII (120 copies) on the inner one (1-4). Inside the capsid, the ~36 kbp linear dsDNA genome is tightly packed together with ~25 MDa of protein: ~500-800 copies of polypeptide VII, ~100-300 copies of polypeptide µ, and 150 copies of polypeptide V (3, 4). Because the three core proteins have a large proportion of positive charges, it is expected that their main role is to help screen the electrostatic repulsion of the confined, negatively charged genome (5).

During maturation, the adenovirus protease (AVP) cleaves the precursor forms of minor coat proteins IIIa, VI, and VIII, as well as core proteins VII and µ, the terminal protein, and the packaging protein L1 52/55 kDa (6-8). AVP enters the viral particle bound to the DNA molecule (its first cofactor) and cleaves 11 residues from the C-terminus of the polypeptide VI precursor (pVI). The released peptide (pVI_C_) is AVP second cofactor, which the protease uses as a sled to scan along the dsDNA and reach the rest of its targets (9-11). These include, among others, a second cleavage site at the N-terminal region of pVI (releasing a 33 residue-long peptide, pVI_N_), and two N-terminal cleavage sites in the precursor of the major core protein, pVII, releasing peptides pVII_N1_ (residues 1-13) and pVII_N2_ (14-24) (6).

Protein VI is located beneath the hexon shell (2, 12) and its membrane lytic activity is crucial for adenovirus cell entry (13-17). Immature adenovirus particles contain the precursors of all viral AVP substrates. They assemble and package correctly, but fail to expose the pVI lytic peptide (an amphipathic α-helix at the N-terminus of protein VI) during entry, become trapped in endo-lysosomes and therefore are not infectious (18-21). This happens because immature particles are more stable than the mature virions (22-24), and do not respond to the cues from receptor movements and confinements when bound to the plasma membrane (16, 25-27). Maturation of the core proteins increases the internal pressure generated by electrostatic repulsion between the confined DNA strands (28). This pressurization seems to provide a mechanism to facilitate penton release upon receptor interaction, which in turn opens the way for exposure of the lytic peptide (22, 23).

Two recent studies have shown unexpected connections between proteins VI and VII. In the first one (29), loxP sites were inserted in the human HAdV-C5 genome flanking the pVII gene (Ad5-VII-loxP). When Ad5-VII-loxP infects cells expressing the Cre-lox recombinase, pVII is not expressed because the VII locus is excised from the viral genome. The first surprising result was that the absence of the major core protein did not hinder particle formation and genome encapsidation. Therefore, the DNA condensation activity of protein VII is not necessary for the adenovirus genome to fit inside the capsid. The second surprise in this study was that, in the particles lacking protein VII (Ad5-VII-), protein VI was present in an intermediate maturation form (iVI), having been cleaved only at its C-terminal site, but not at the N-terminus. Additionally, Ostapchuk and collaborators found that the Ad5-VII-particles were not infectious due to a defect in entry: they were unable to escape from the endosome, similarly to the fully immature particles produced by the thermosensitive mutant Ad2 *ts1* (19, 30).

Why full maturation of pVI is impaired in the absence of core protein VII, and why the particles lacking VII are trapped in the endosome, has remained unclear. The fact that pVI was processed at the C-terminus means that AVP was able to produce its cofactor (pVI_C_), become activated, and process the rest of its substrates, because neither full length L1 52/55 kDa nor pVIII were detected in Ad5-VII-particles (29). However, AVP could not cleave pVI_N_, which *in vitro* is the first site processed in pVI (10). The N-terminal cleavage of pVI occurs directly upstream of the region containing the amphipathic α-helix (residues 34-54) involved in endosome membrane disruption (13, 14). This cleavage may not be required for the lytic activity of the helix, because recombinant pVI is able to disrupt liposome membranes *in vitro* (13).

The second study revealing unexpected relationships between proteins VI and VII is the most recent cryo-electron microscopy (cryo-EM) analysis of the HAdV-C5 capsid structure (2). Structural and biochemical analyses had previously shown that the pVI_N_ peptide is located inside a central cavity of the hexon trimer that opens towards the viral core, but density for pVI_N_ was weak and present only in a few hexons, indicating lack of icosahedral order, or partial occupancy (12, 31, 32). Partial occupancy is also implied by the hexon and pVI copy numbers in the virion: there are too many copies of pVI (~360) to have one per hexon trimer (240), but too few to have one per hexon monomer (720). In (2), an increase from 3.5 to 3.2 Å resolution provided new information on the molecules generating weak density inside hexons. First, at the mouth of the hexon cavity, residues 109 to 143 of pVI were modeled, disconnected from pVI_N_. Second, the pVI_N_ tracing was reversed with respect to the previous structures (31, 32), in such a way that in the latest model (2) the pVI_N_ cleavage site is not accessible at the rim of the hexon cavity, but hidden inside and oriented away from the core. This new disposition makes it more difficult to understand how AVP, sliding on the DNA, can reach its target sequence in pVI_N_ (**Fig. S1**). Third, it was shown that one of the densities inside the hexon cavity did not correspond to pVI_N_, but to the second N-terminal peptide cleaved from core protein VII (pVII_N2_). This observation was also unexpected, because as a core protein, no part of VII was thought to be icosahedrally ordered, and there were no previous indications that pVII interacted with hexon. The pVII_N2_ peptide occupies a position equivalent to that of pVI_N_, lining the hexon cavity wall. Residues Ser31 in pVI_N_ and Phe22 of pVII_N2_ interact with the same binding pocket in the hexon wall, raising the possibility that pVI and pVII may compete for hexon binding during assembly (2).

In another report, we have analyzed how the absence of protein VII impinges on the mechanical properties and core organization of the HAdV-C5 particle (33). We showed that the entry defect of Ad5-VII-particles could not be attributed to either lack of internal pressure or excessive binding of the naked DNA to the capsid shell. Here, we analyze exposure of protein VI, capsid stability and capsid structure in Ad5-VII-, in search of the reasons why these particles are impaired in pVI maturation and endosomal escape. Our results show for the first time the organization of the pVI lytic peptide inside the hexon cavity before pVI_N_ cleavage, and provide the basis for a model in which competition between the precursors pVI and pVII for hexon binding is crucial to regulate adenovirus maturation, and therefore infectivity.

## Results

### Exposure of membrane lytic protein VI is impaired in the absence of core protein VII

For protein VI to mediate endosome disruption, the lytic peptide (amphipathic helix in residues 34-54) needs to be released from the interior of the viral particle (13). Therefore, we investigated whether lack of protein VII had an effect on exposure of protein VI upon viral entry. To this end, we characterized the exposure of protein VI in human A549 cells infected with atto 565-labelled viral particles upon cell entry. The signal for protein VI in HAdV-C5 wildtype particles (Ad5-wt) internalized particles increased between 5 and 10 times during the first 10 min of infection, indicating progressive exposure of VI from the incoming particles (Fig. 1). This was consistent with earlier reports demonstrating exposure and complete loss of VI from the virions upon endosomal rupture and virion release to the cytosol (16, 34). In simultaneous experiments with Ad5-wt and Ad5-VII-particles (Fig. 1), we observed that the Ad5-VII-virus tended to show higher amounts of protein VI signal at the 0 min time point than the Ad5-wt virus, suggesting the presence of damaged particles in the preparations, a possibility that was supported by negative staining electron microscopy (EM) imaging (**Fig. S2).** Unlike Ad5-wt, Ad5-VII- did not show an increase in protein VI exposure in the first 20 min of infection after warming of cold synchronized cells (Fig. 1b). Nevertheless, the Ad5-VII-virus did not present difficulties in cell internalization (**Fig. S3**). These results suggest that Ad5-VII-preparations contain some damaged particles, which likely do not internalize, and the reminder of the Ad5-VII-particles fail to properly expose the partially processed polypeptide VI as they enter cells. This observation is consistent with the reported failure of Ad5-VII-to penetrate the endosome membrane (29).

**Figure 1.**
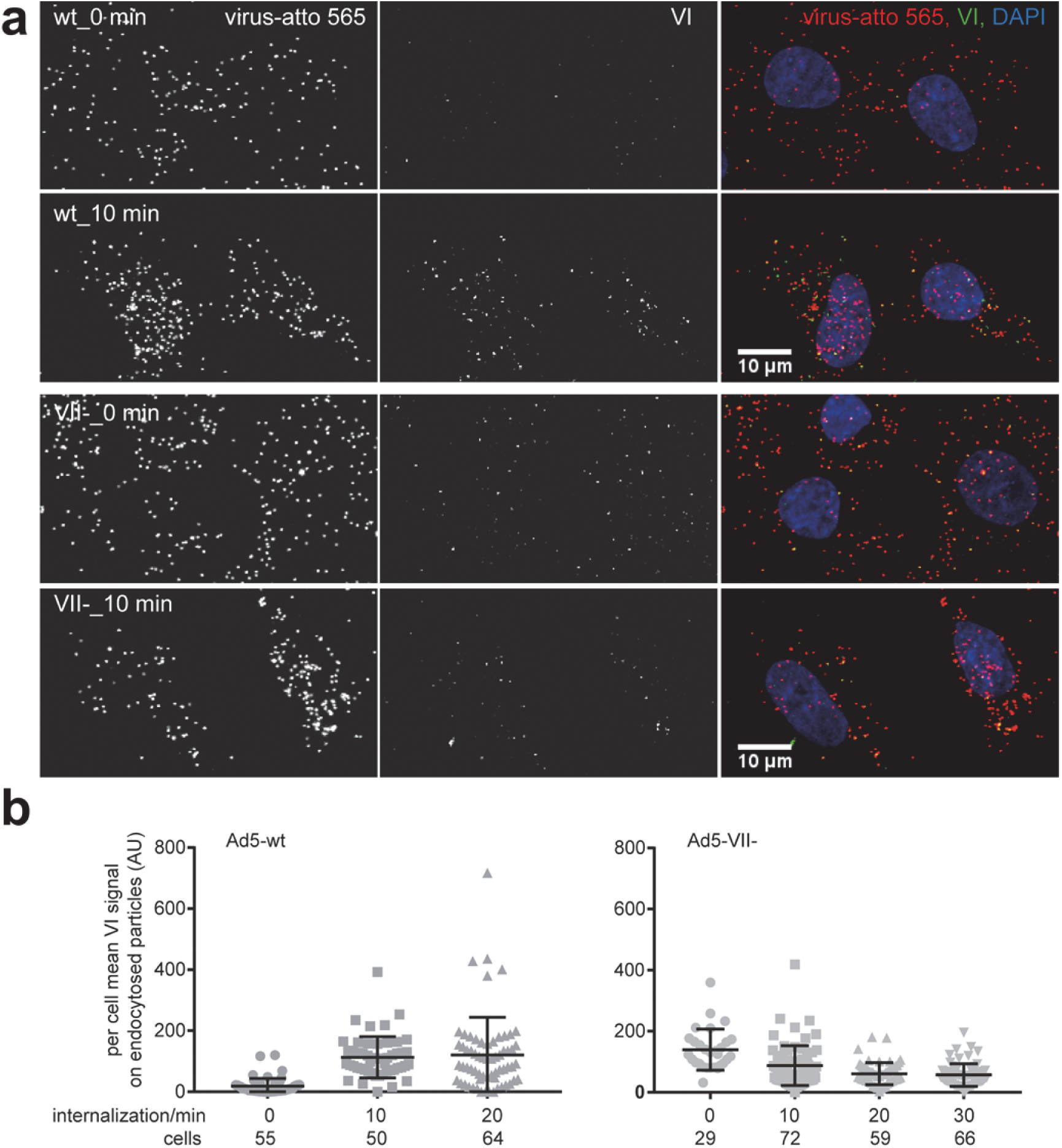
Reduced protein VI exposure in Ad5-VII-particles during entry in comparison to Ad5-wt. Atto 565-labeled virus particles were added to A549 cells at 4 °C for 60 min. Unbound particles were washed away, and cells were shifted to 37 °C for the indicated times. Intact cells were then incubated with mouse 9C12 anti-hexon antibody at 4 ºC to tag plasma membrane-associated viruses, and after fixation, cells were permeabilized and stained with rabbit anti-protein VI antibody. These primary antibodies were detected with Alexa Fluor 680-conjugated anti-mouse and Alexa Fluor 488-conjugated anti-rabbit antibodies. Nuclei were stained with DAPI, and samples were imaged by confocal microscopy. **(a)** Representative images shown for Ad5-wt (top rows) and Ad5-VII-(bottom rows) at the 0 min and 10 min time points are maximum projections of confocal stacks. Scale bars = 10 µm. **(b)** The scatter plots show per cell mean protein VI signal on endocytosed particles (i.e., particles devoid of 9C12 antibody signal) for 10 min and 20 min time points, but the 0 min time point includes all cell-associated particles. One dot represents one cell and the number of cells analyzed is indicated. Error bars represent the means ± SD.

### Adenovirus particles lacking protein VII have reduced thermal and mechanical stability

The Ad5-VII-entry defect is reminiscent of that observed for particles produced by the Ad2 *ts1* mutant, which contain immature pVI and become trapped in the endocytic route (19, 30, 35). Ad2 *ts1* particles are highly stable: they can withstand harsher conditions than Ad5-wt, require more stress to release pentons, and therefore do not release protein VI upon entry (16, 18, 19, 22-24). Likewise, mutation G33A in pVI produces particles with partial processing of this protein, higher thermostability, lower membrane lytic activity, reduced infectivity, and more stable pentons (36, 37). To evaluate whether increased stability could be responsible for the lack of protein VI release from Ad5-VII-particles, we carried out thermal and mechanical disruption assays.

We used extrinsic fluorescence of the DNA intercalating agent YOYO-1 to characterize capsid disruption as a function of temperature in Ad5-wt and A5-VII-particles. In both cases, the fluorescence emission of YOYO-1 increased with the temperature, as the genome became exposed to the solvent (Fig. 2). The half-transition temperature (T_0.5_) estimated for Ad5-wt was 47.35 ± 0.06 ºC, in agreement with previous studies (13, 22, 24, 38, 39). In contrast, for Ad5-VII-the maximum rate of DNA exposure happened at 44.45 ± 0.10 ºC. This result indicates that Ad5-VII-particles are thermally less stable than Ad5-wt.

**Figure 2.**
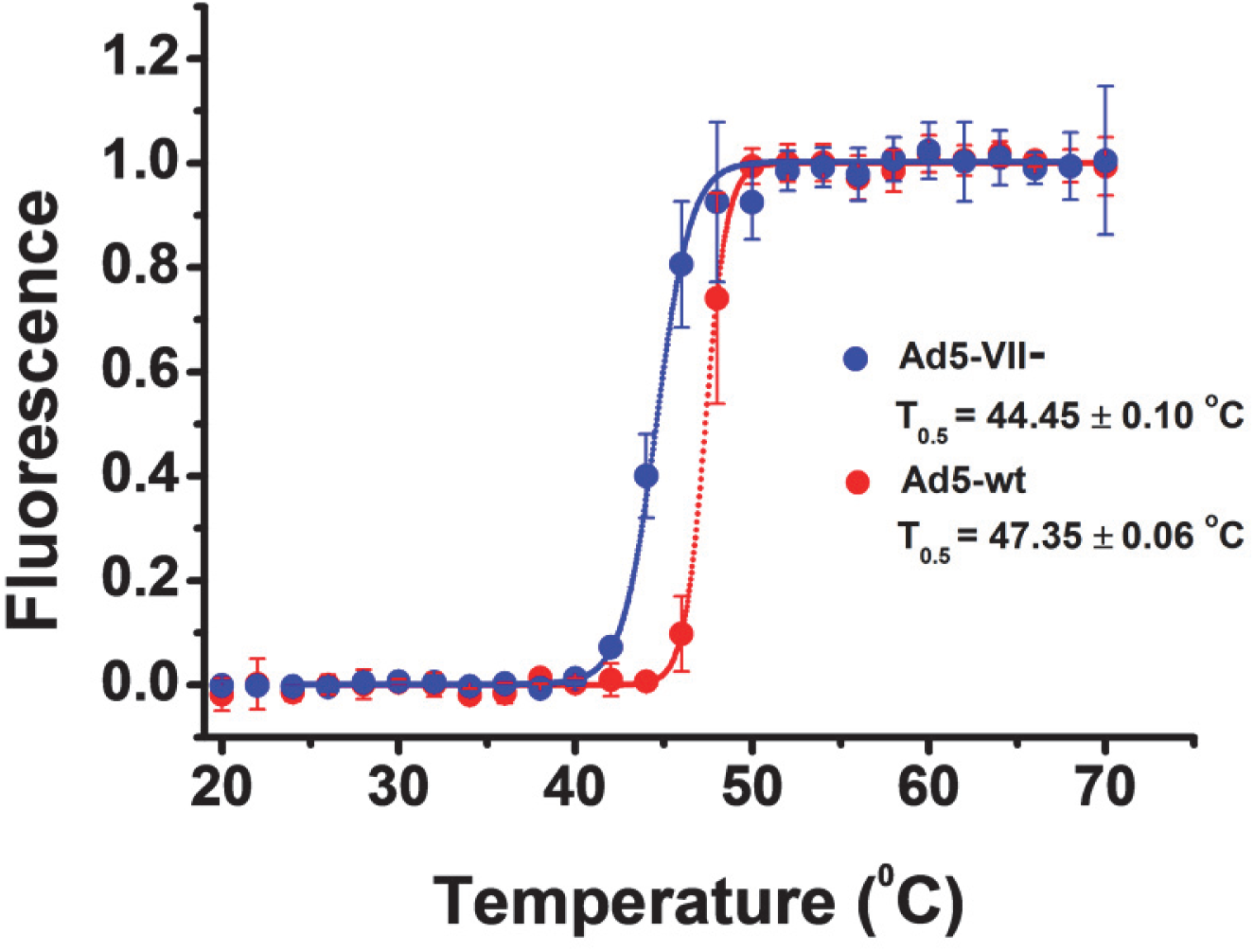
Thermal stability of adenovirus particles with or without protein VII. Extrinsic fluorescence measurements showing DNA exposure to solvent upon temperature increase for Ad5-VII-(blue) and Ad5-wt (red). Average values and *error bars* corresponding to the standard deviation of four experiments are plotted. Dotted lines correspond to the fit according to a Boltzmann sigmoid function.

To analyze the mechanical stability of the Ad5-VII-particles, we used mechanical fatigue induced by atomic force microscopy (AFM). This method involves repeated imaging of the same virus particle using very low forces (~100 pN), far below the breaking force of the capsid. In this way, we simultaneously induce and monitor the gradual disassembly of single virus particles (23). Topography images taken at the beginning of the experiment showed no differences between Ad5-wt and Ad5-VII-particles (Fig. 3a, **panels #1**). As the experiment proceeded, mechanical fatigue caused the sequential removal of pentons, followed by complete particle collapse. In the example shown in Fig. 3a, we show that for Ad5-VII-the first penton fell off the capsid at image #3, and the third penton at image #9. The capsid collapsed after #18 images. In contrast, in the Ad5-wt the first penton falls off at image #43, and the capsid collapsed at image #59.

**Figure 3.**
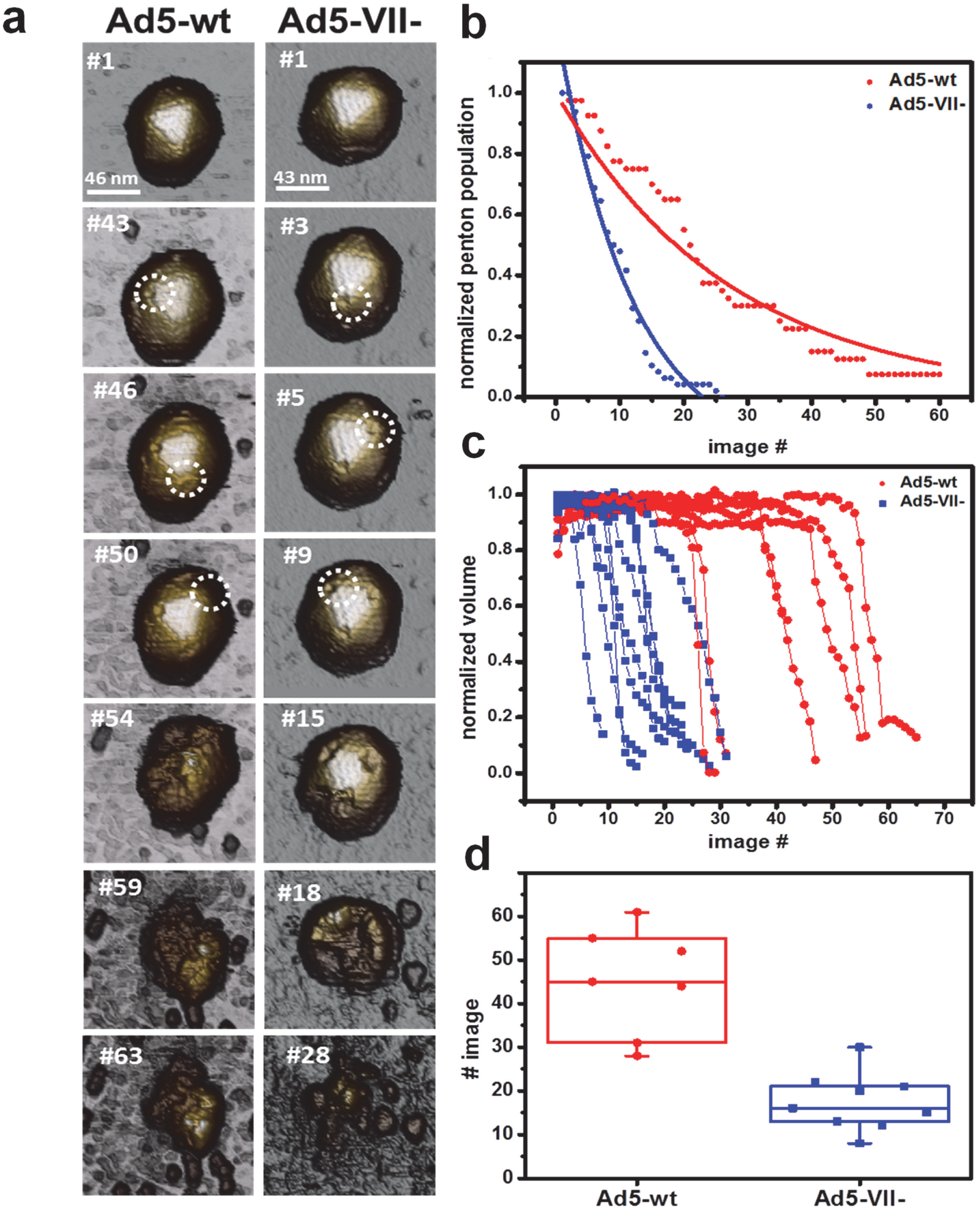
Mechanical stability of adenovirus particles with or without protein VII. **(a)** Representative AFM images along fatigue experiments for Ad5–VII-and Ad5-wt particles adsorbed on mica. Release of the first, second and third pentons from the top facet are indicated with white circles. **(b)** Analysis of penton release for 9 Ad5-VII-and 7 Ad5-wt viral particles. At the beginning of the experiment, all visible pentons in the images are occupied (penton population = 1). The penton population decreases along the experiment. **(c)** Comparison of the evolution of Ad5-VII-and Ad5-wt particle volume during the fatigue experiment. **(d)** The image number at which each particle has undergone a 50% volume decrease is plotted for all particles analyzed.

We quantified the rate of penton loss for 9 Ad5-VII-and 7 Ad5-wt viral particles. In Fig. 3b, pentons are considered independent identities, and the normalized penton population for all studied particles is plotted along the number of images taken. At the beginning of the experiment, all observable pentons are present for the 16 studied particles (penton population = 1). As the experiment proceeds, pentons fall off and the penton population decreases until there are no more observable pentons. The penton release distribution follows an exponential decay for both types of particles. The mean life-time of a penton is 12 images for Ad5-VII-, and 26 images for Ad5-wt, indicating that penton loss occurs faster in Ad5-VII-particles (Table 1). This higher propensity to release pentons agrees with our observations showing that lack of core protein VII increases the particle internal pressure(33).

**Table 1.** Exponential decay distribution of penton release. The penton distribution data follow an exponential decay function N(t)=N_0_e^−λt^, where *t* is the number of images taken, and parameters such as the penton release rate constant (λ) and mean life-time of a penton (1/λ) can be determined. The mean penton lifetime is given as number of images; the penton release rate constant represents the probability to release one penton in one image.

| Specimen | Penton release constant $\lambda$ (images<sup>-1</sup>) | Mean penton lifetime $1/\lambda$ (images) | N |
| --- | --- | --- | --- |
| Ad5-VII- | 0.08 | 12 | 9 |
| Ad5-wt | 0.04 | 26 | 7 |

To further analyze the disassembly dynamics, we quantified the variation in particle volume during the mechanical fatigue assays (Fig. 3c). At the beginning of the experiment, the particle volume remained constant for all particles analyzed. After losing the three pentons on the upper facet, the volume drops at a similar rate for both kinds of particles (13% volume loss/image for Ad5-VII-, 14% volume loss/image for Ad5-wt). A clear difference, however, was the time elapsed until the particle volume starts to decrease: Ad5-wt particles lost 50% of their volume after being imaged ~50 times, while for Ad5-VII-the same decrease in volume occurred after only 15 images (Fig. 3d). That is, Ad5-VII-particles start losing volume earlier than Ad5-wt particles, consistent with the penton release tendency. Therefore, penton resilience determines the stability of the virus particle under mechanical fatigue.

In conclusion, our results so far indicate that Ad5-VII-particles are less stable than Ad5-wt under both thermal and mechanical stress. Therefore, the defect of Ad5-VII-virions in protein VI exposure is unlikely due to higher mechanical or thermal stability, although it might be due to an inability of the VII-virions to respond to the pulling forces from CAR receptor movements and holding forces of the immobile integrins.

### Lack of density in the Ad5-VII-cryo-EM map hexon cavity confirms the interaction of core protein VII with hexon

Since the absence of protein VII decreases the stability of the HAdV-C5 capsid, it would be expected that exposure of protein VI was easier in Ad5-VII-than in Ad5-wt. However, this is not the case. To try to understand why, we investigated the structural differences between Ad5-VII- and Ad5-wt particles using cryo-electron microscopy (cryo-EM) (**Fig. S4**). A difference map was calculated by subtracting the Ad5-VII-map from Ad5-wt, to show density corresponding to elements present in the mature virion but absent when the major core protein is missing. This map (Fig. 4a, red density) showed small differences located in the interior of the capsid, near the rim of the hexon cavity (red circles, Fig. 4a). When we overlap the most recent Ad5-wt structure (PDB-6B1T (2)) with our maps, we observe that one of the small densities absent in Ad5-VII-perfectly matches the position of peptide VII_N2_ traced by Dai et al. inside the cavity of hexon 2 in the Asymmetric Unit (AU) (Fig. 4b and 4c).

**Figure 4.**
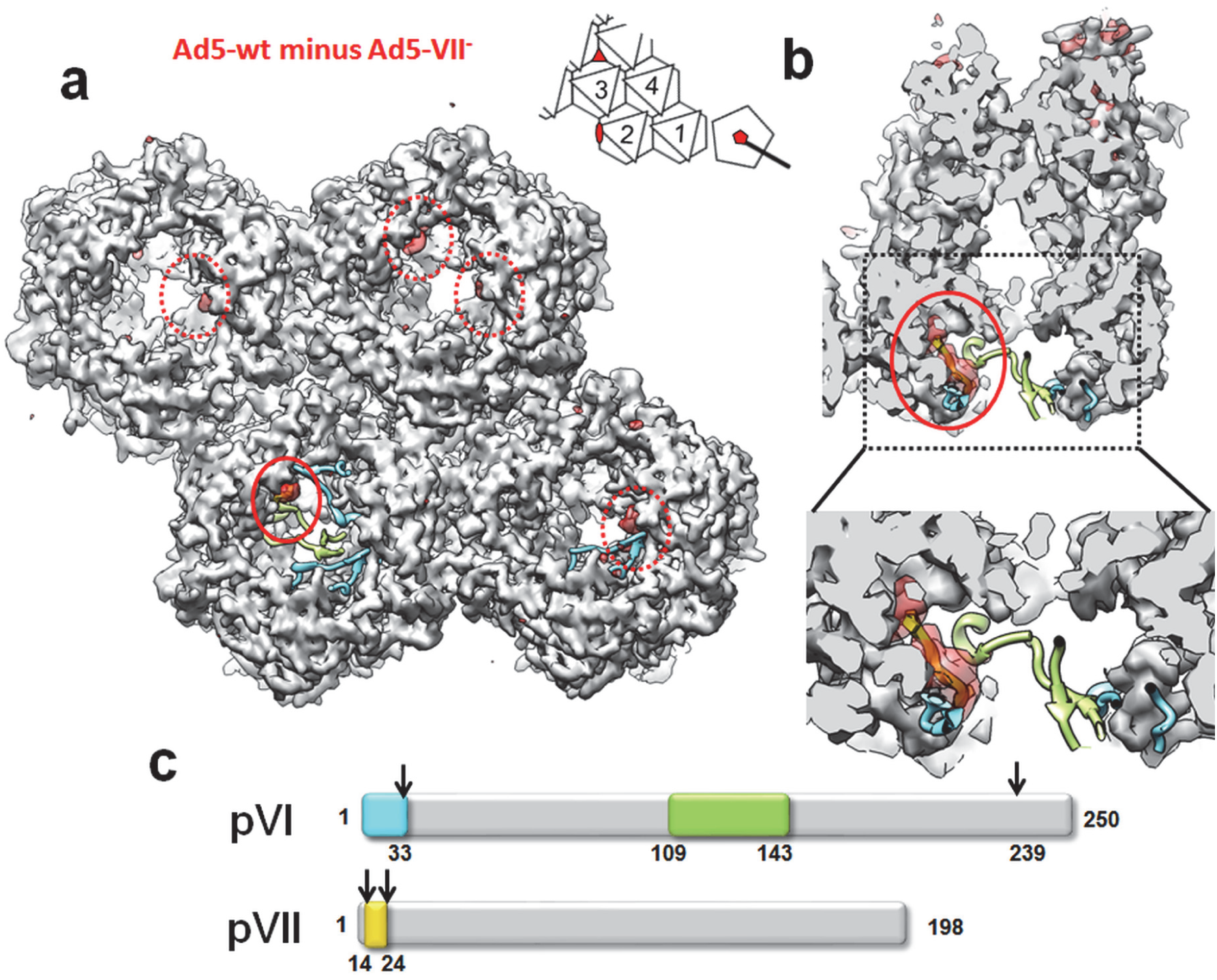
Cryo-EM difference map showing density present in Ad5-wt but absent in Ad5-VII-particles. **(a)** The four hexon trimers in the AU (see inset) as seen from inside the viral particle are shown as a gray surface generated from the high resolution HAdV-C5 structure (PDB 6B1T). The difference map obtained by subtracting the two cryo-EM maps (Ad5-wt minus Ad5-VII-) is shown as a semitransparent red surface contoured at 1.5σ level. Cyan, green and yellow ribbons correspond to the pVI and pVII fragments traced in (2), as indicated in the schematics in (c). The solid line red circle indicates the location of the pVII_N2_ peptide (yellow ribbon, covered by the semitransparent difference map) traced by Dai et al. The dashed red circles indicate the position of four additional difference densities located at positions in the hexon cavity where either pVI_N_ or pVII_N2_ would bind, according to Dai et al. To remove visual clutter, only the 150 largest difference patches within a radius of 15 Å around the four hexon trimers are shown (UCSF Chimera *sop zone # selected 15 maxComponents 150*). The inset shows a schematic of the icosahedral asymmetric unit as seen from inside the particle, with the four hexon trimers numbered 1-4, and the positions of the 2, 3 and 5-fold icosahedral symmetry axes depicted by a red oval, triangle and pentagon, respectively. **(b)** Cutout view of hexon 2 showing that the difference density clearly fits with the position of pVII_N2_ modeled by Dai et al. **(c)** Schematic representation of proteins pVI and pVII. Numbers at the right hand side indicate the length of each protein (in amino acids). Arrows point to the AVP cleavage sites. Colored boxes represent the fragments solved by Dai et al.

When the difference map is contoured at 1.5σ, we observe five red densities in the AU, one inside each of hexon trimers 1, 2 and 3, and two inside hexon 4. These difference densities occupy positions near the VI_N_/VII_N2_ binding pocket in hexon, according to (2). Since there are ~360 copies of pVI (3, 4) and 720 hexon monomers (organized in 240 trimers) in the capsid, at least half the possible binding pockets (six out of twelve per AU) would be available for pVII to bind, in good agreement with the observation of 5 empty pockets when VII is not present (Fig. 4a). However, all difference densities are weaker than the density corresponding to hexon (3σ), indicating disorder or partial occupancy. At lower contour level (1σ), 8 red densities located near the VI_N_/VII_N2_ binding pocket in hexon can be observed in the difference map AU (**Fig. S5**). One of these, located in the peripentonal hexon (hexon 1) cavity, covers the space occupied by one of the pVI_N_ peptides traced in the Ad5-wt structure. One more difference blob per AU appears at nearly noise level (0.8σ, **Fig. S5**).

Given that protein VII is the only component missing from the Ad5-VII-particle (29), we interpret the (Ad5-wt minus Ad5-VII-) difference densities as an indication of binding sites left empty by the absence of the major core protein. This result supports the location of part of protein VII (the pVII_N2_ peptide) proposed by the latest cryo-EM study on Ad5-wt (2). The fact that more difference blobs appear in equivalent capsid positions at lower contour thresholds is consistent with variable occupancy of the pVI_N_/pVII_N2_ binding pockets across the different hexon monomers icosahedrally averaged in the cryo-EM map. This result indicates that in the complete virion, the pVII_N2_ peptide has the potential to bind all hexon monomers in the capsid, but succeeds only in part of them, while in the rest the binding pocket is occupied by pVI_N_. That is, our observations are consistent with the hypothesis that during Ad5 assembly, pVI and pVII compete for the same binding site in the hexon trimer cavity.

### Observation of partially processed protein VI in the Ad5-VII-particle

The inverse difference map showed the density corresponding to icosahedrally ordered elements present in Ad5-VII-but absent in Ad5-wt. Direct observation of the cryo-EM maps already revealed a clear density filling the inner cavity of hexons in Ad5-VII- but absent in Ad5-wt (**Fig. S4**, white circle). This extra density is present in all hexons in the AU (Fig. 5 **and Fig. S6**), and is consistent with an increase in order for unprocessed polypeptide VI, as shown previously for the immature particle (Ad2 *ts1*) (24, 40). Even when contoured at low density (0.8σ, Fig. 5), the negative difference does not cover the pVI fragments traced in the latest Ad5-wt model (neither pVI_N_ nor the central VI fragment), indicating that these regions are mostly organized in the same way in both Ad5-wt and Ad5-VII-.

**Figure 5.**
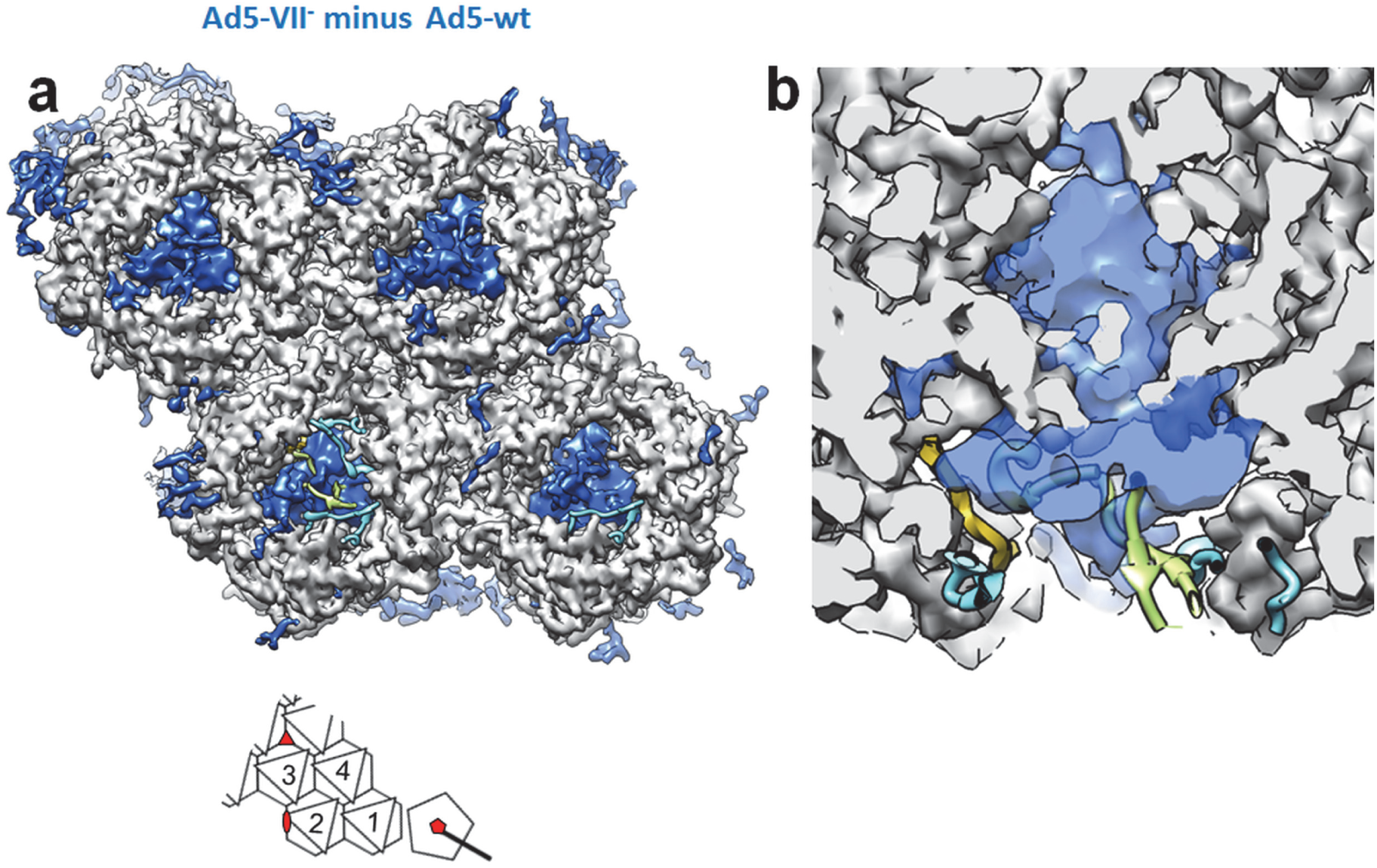
Cryo-EM difference map showing density present in Ad5-VII- but absent in Ad5-wt particles. **(a)** The four hexon trimers in the AU (see inset) as seen from inside the viral particle are shown as a gray surface generated from the HAdV-C5 high resolution structure (PDB 6B1T). The difference map obtained by subtracting the two cryo-EM maps (Ad5-VII-minus Ad5-wt) is shown as a blue surface and contoured at 0.8σ level. Cyan, green and yellow ribbons correspond to the pVI and pVII fragments traced by (2), as in Fig. 4. To remove visual clutter, only the 80 largest difference patches within a radius of 15 Å around the four hexon trimers are shown (UCSF Chimera *sop zone # selected 15 maxComponents 80*). **(b)** Zoom in on a cutout view of the hexon 2 trimer showing that even at this low contouring threshold the difference density does not cover the pVI and pVII fragments modeled by Dai et al.

When, instead of calculating the difference map between the two cryo-EM maps, we subtract only the hexon density (generated from the high resolution structure) from the Ad5-VII-map, the conformation of the extra density inside the hexon cavity can be analyzed in more detail. In hexon 2, this density forms two well defined, L-shaped tubular regions directly connected to the C-terminal residue of the pVI_N_ peptides traced by (2) (Fig. 6a). Secondary structure predictions and circular dichroism analyses indicate that the N-terminal region of protein VI after cleavage by AVP is predominantly α-helical (**Fig. S7**) (14). One of these helices would harbor the amphipathic, lytic peptide (Ala34-Tyr54) (13). Indeed, two α-helices, 11 and 18 residues long respectively, can be modeled in each of the two L-shaped densities inside the hexon cavity (Fig. 6b). Together with the continuity with the pVI_N_ peptides traced in the high-resolution structure, the shape and size of the difference density indicate that each one of the two L-shaped tubes corresponds to ~30 residues of protein VI, still uncleaved from the pVI_N_ peptide. From the last residue in these helices (~residue 65), a ~50 residue-long disordered region would connect with the central fragment of the protein traced in (2), which starts at residue 109 (Fig 6c). Although at a limited resolution which precludes identification of side chains, this is the first observation of the adenovirus lytic peptide organization within the capsid.

**Figure 6.**
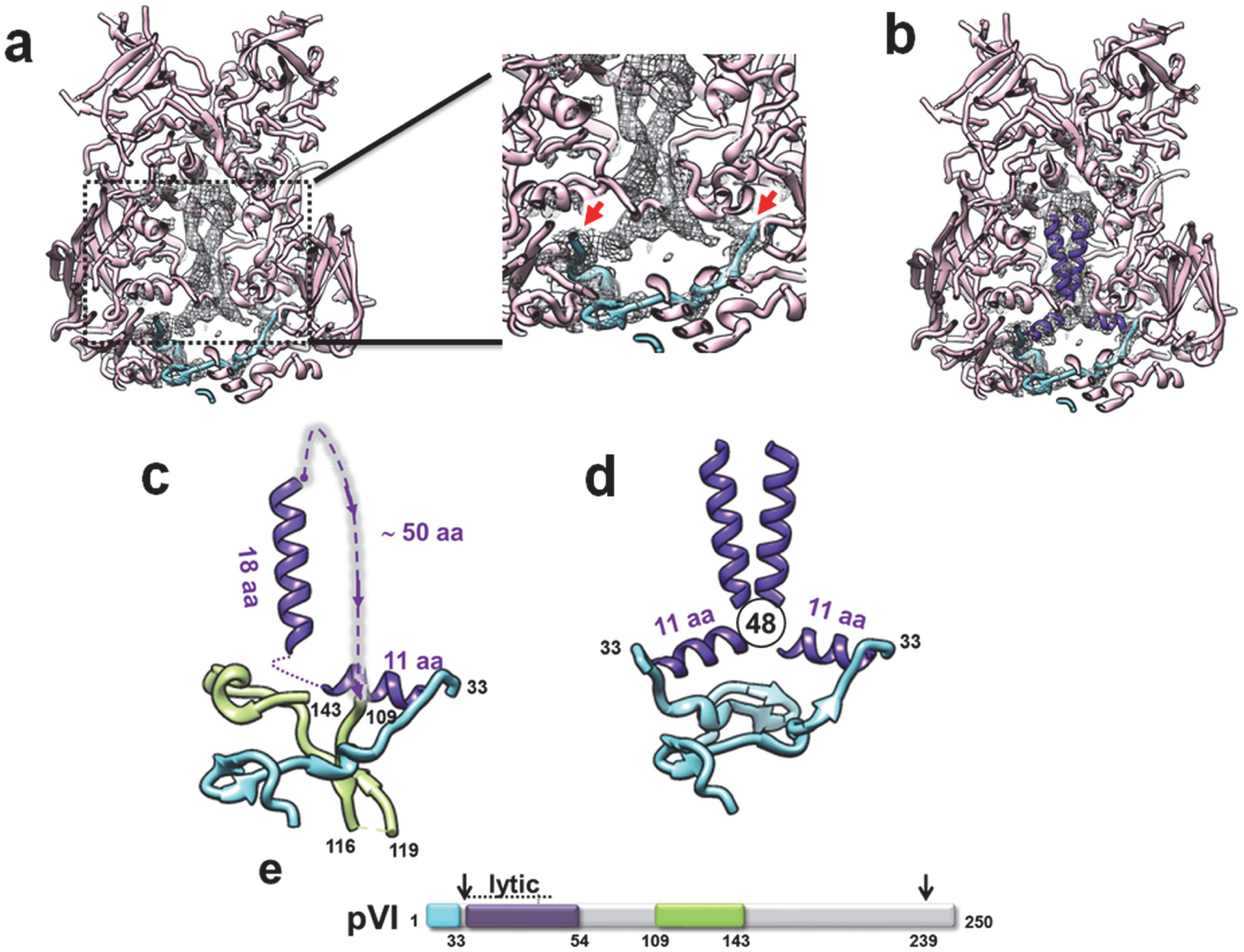
Ordered density inside the hexon cavity in Ad5-VII-. **(a)** A difference map obtained by subtracting the hexon 2 density (calculated at 5.5 Å resolution from PDB 6B1T) from the Ad5-VII-cryo-EM map, and contoured at 1.7σ, is shown as a gray mesh. Hexon 2 (pink) and two copies of the pVI_N_ peptide (cyan) as traced in PDB 6B1T are represented as ribbons. The zoom at the right hand side shows the continuity between the difference density and the pVI_N_ peptides traced by Dai et al. (red arrows) **(b)** As in (a), with two α-helices (11 and 18 residue long, respectively) modeled in each L-shaped tubular density inside the hexon cavity (purple). For a cleaner view, only a region of the difference map contained within 12 Å from the modeled helices is shown. **(c)** Model for the organization of one pVI monomer in the virion. The pVI_N_ peptide traced by Dai et al. (cyan) is followed by the two α-helices modeled in the Ad5-VII-map (purple). A dotted line indicates the connection between helices, and the dashed line indicates a disordered region connecting the end of the modeled helices to the central region of pVI as traced by Dai et al. The length of each helix and of the disordered segment is indicated (in amino acids). Residue numbers in black correspond to the limits of the pVI regions traced by Dai et al. **(d)** Model for the pVI dimer showing how our model places residue 48 of the two monomers in the hexon cavity in close proximity. **(e)** Schematic representation of polypeptide pVI as in Fig. 4c, with the position of the pVI lytic peptide indicated as a purple box.

At the same contouring threshold, we also observed two L-shaped tubular densities inside the hexon 3 cavity, while in hexons 1 and 4 the density for the second tube was weaker (**Fig. S8**). These results contribute to explain the long standing puzzle of the stoichiometry of polypeptide VI. Given the available space, it seems very difficult to fit one copy of VI per hexon monomer inside the trimer cavity. Hexons 2 and 3 seem to have a preference for accomodating two copies of VI, while in hexons 1 and 4, although two copies can be held inside the cavity, there is a greater probability of finding only one. The sum of 2+2+1+1 copies of VI will result in the previously estimated number of 6 copies per AU. Interestingly, hexons 1 and 4 differ from hexons 2 and 3 in their interactions with another minor coat protein, polypeptide VIII (1), suggesting that the presence of this minor coat protein plays a role in determining the hexon:VI stoichiometry. Our interpretation of the densities in the hexon cavity is also consistent with the observation of dimeric species when a cysteine was introduced at position 48 in pVI (G48C, (41). In our model, the region where residue 48 would be located corresponds to the elbow of the L-shaped density tubes, placing the two cysteines (one from each pVI molecule) close enough for a disulfide bond to form (Fig. 6d). Dimers of protein VI were also detected by crosslinking assays on disrupted virions and extracted cores (42)

It has previously been shown that immature pVI has lytic activity, but binding to hexon shields this activity (36). The mature form of VI binds to the hexon cavity more weakly than the intermediate one (iVI), as indicated by assays where heating of Ad5 particles carrying the pVI G33A mutation caused release of mature protein VI, but not of iVI (36). Our difference mapping is in agreement with these observations: when core protein VII is not present and the N-terminal cleavage of VI does not occur, the lytic peptide remains shielded by hexon, even if capsid disassembly proceeded unimpeded. Alternatively, it is possible that the capsid of Ad5-VII-particles does not disassemble in cells, which would be supported by the observation that the incoming viral DNA in Ad5-VII-particles is less accessible to click chemistry labeling than the Ad5-wt DNA (29).

## Discussion

When Ad5 protein VII is not present during assembly, pVI_N_ is not cleaved, and the viral particle remains trapped in the endosome (29). The endosome escape defect is unlikely due to the inability of pVI_N_ to lyse membranes, as shown by *in vitro* experiments with recombinant pVI (13). Rather, what happens is that in Ad5-VII-particles, protein VI is not released in the endosome (Fig. 1), preventing the interaction between its amphipathic helix and the endosome membrane. The inability to release protein VI is not due to increased physical stability of the Ad5-VII-particles (Fig. 2 and 3). Rather, the lytic peptide remains fused to the pVI_N_ peptide, and is hidden within the hexon cavity, unavailable to interact with the endosome membrane (Fig. 6).

One intriguing question is prominent in the analysis of the Ad5-VII-particles: why is pVI_N_ not cleaved when protein VII is absent? Indeed, it is not clear yet how pVI_N_ is cleaved during assembly in Ad5-wt, because the highest resolution structure reported so far indicates that the pVI_N_ peptide is oriented with its cleavage site hidden inside the hexon cavity, away from the reach of the DNA-bound AVP (**Fig. S1**) (2). Our observations provide new clues to answer these questions, prompting a new model for the role of protein VII in regulating capsid maturation.

Dai et al interpreted part of the density inside the cavity of one of the hexons in the AU as a peptide cleaved not from pVI, but from pVII, suggesting that the two proteins could compete for the same binding site in hexon during assembly (2). Our cryo-EM difference maps indicate that pVII_N2_ has the ability to bind to the same pocket as pVI_N_ in all 12 hexon monomers in the AU. In one viral particle, there are 500-800 copies of pVII, and 360 copies of pVI (3, 4). That is, there are 860-1160 candidates to bind to 720 sites in the hexon trimer cavities. We propose that, due to this excess, proteins VI and VII establish a dynamic competition for hexon binding sites during assembly. As a result of this competition, pVI or pVII molecules carrying their hexon-binding N-terminal peptides would continually be pushed out from the hexon cavity by their competitors, putting them in the path of AVP sliding on the DNA (Fig. 7). In the absence of protein VII, the competition does not exist: there are enough sites to bind for all pVI copies, and nothing to push the lytic peptide out from the hexon cavity. Thereby, the pVI_N_ cleavage site would be hidden away from the protease. Consequently, (a) pVI_N_ is not cleaved, and (b) the lytic peptide remains shielded by hexon, preventing virion escape from the endosome.

**Figure 7:**
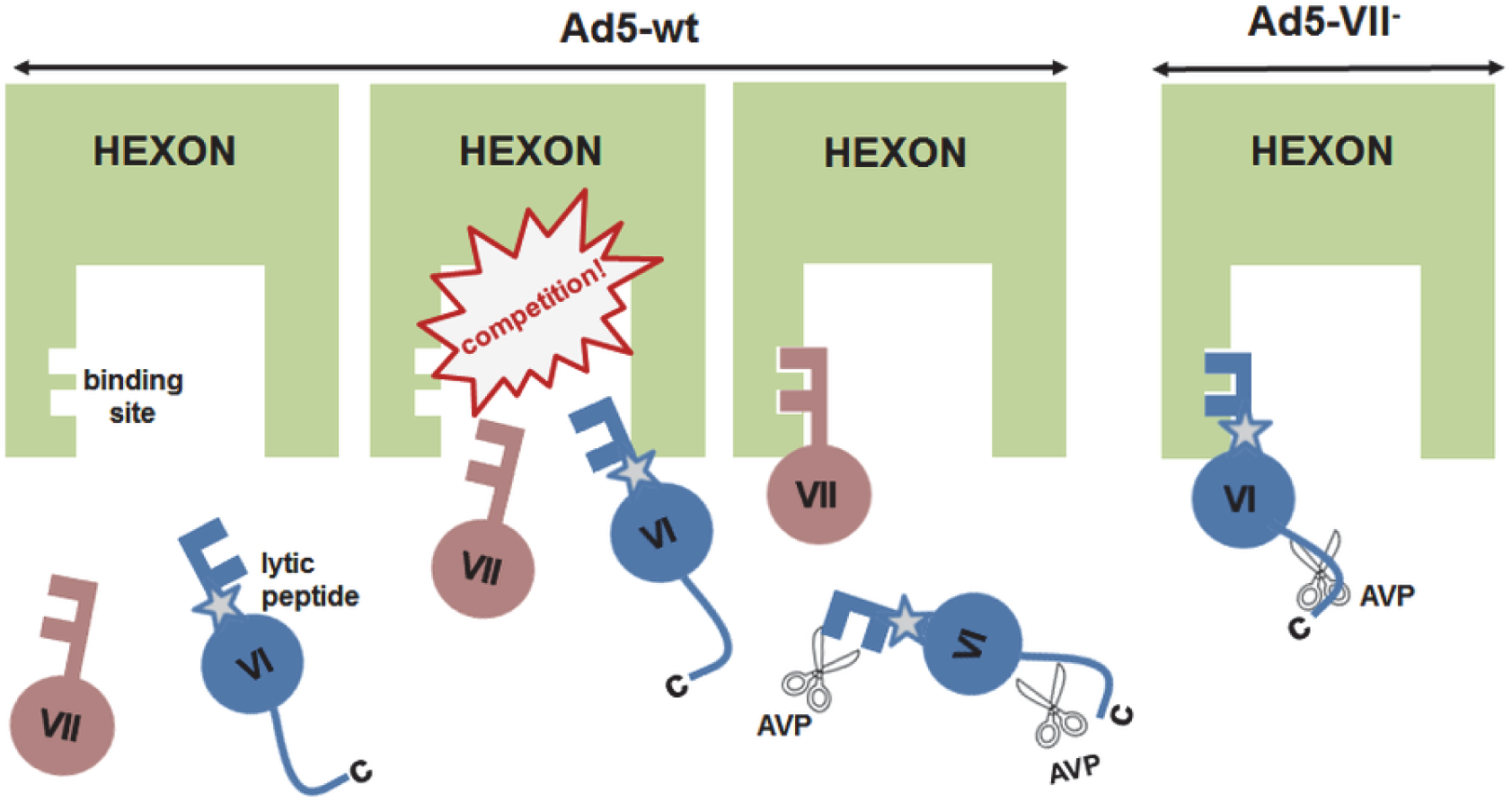
Model for dynamic competition between polypeptides VI and VII. During Ad5-wt assembly, the N-terminal regions of pVI and pVII (keys) compete for the same binding pocket in the hexon monomer (lock). This competition forces one of the two binding candidates out of the hexon cavity, making its N-terminal cleavage site accessible to AVP (scissors), and in the case of protein VI, for exposure and membrane disruption in the endosome. In Ad5-VII-, pVI has no competitors, and therefore its N-terminal region remains bound to and hidden inside the hexon cavity, unavailable for proteolytic cleavage or membrane disruption.

The competition model we propose here may help explain previous observations on the effect of pVI mutations in adenovirus maturation and infectivity. In mutant G33A, pVI_N_ is not properly cleaved either, but the virion retains a considerable infectious potential (25%) (36). This infectivity can be caused by the presence of 50% properly cleaved VI copies, but our competition model highlights another difference between G33A and Ad5-VII-particles. In G33A, although pVI_N_ is partially uncleaved, the competition with protein VII is present, which could cause the lytic region to be bumped out of the hexon cavity and available for endosome membrane disruption. It has also been observed that in pVI-S28C mutant capsids, pVI was correctly processed at both ends, but a considerable proportion of protein VII remained uncleaved (37). It is possible that the change in a region close to Ser31 in pVI (the residue filling the common pVI/pVII binding pocket in hexon) decreases its affinity for hexon sites in such a way that binding of the more abundant pVII would predominate, with its subsequent protection from the protease. The presence of uncleaved pVII bound to the hexon cavities would result in enhanced bridging between shell and genome, possibly causing the observed stiffening of the S28C particles. In agreement with the correct maturation of pVI, in this mutant infectivity was slightly (2-fold) higher than in the parental virus, indicating that exposure of protein VI during entry was unaltered, or even slightly enhanced.

In the immature particles produced by Ad2 *ts1*, both pVII and pVI are present in their appropriate copy numbers. Therefore, it would be expected that part of the pVI molecules are forced out from the hexon cavity by competition with pVII. Although they would be uncleaved due to the mutation in AVP, in principle these molecules could be exposed and disrupt the endosome membrane. However, the defect in VI exposure in Ad2 *ts1* seems to be related to the lower internal pressure exerted by the genome when condensed by the precursor versions of core proteins VII and µ, resulting in increased capsid stability and delayed penton loss (22, 24, 28, 43).

## Materials and Methods

### Virus specimens

The Ad5-wt virus used in cryo-EM, AFM and extrinsic fluorescence assays was the E1-E3 deleted, GFP expressing Ad5/attP vector which is structurally wildtype (44). The parent virus Ad5-VII-contains loxP sites flanking the protein VII open reading frame and was generated and propagated as described (29). Ad5-VII-particles used in entry assays were produced using A549 cells expressing a Cre recombinase, isolated and labeled with atto 565 as previously described (8, 19, 45). More information can be found in Supplemental Information (SI).

### Endocytosis and protein VI exposure assays

Ad5-wt and Ad5-VII-particles labeled with atto 565 were added to cells at 4 °C for 60 min. Unbound virus particles were removed, cells were washed once with cold medium and incubated in a 37 °C water bath for 0, 10 or 20 min. Afterwards, intact cells were incubated with mouse 9C12 anti-hexon antibody at 4 ºC to tag particles at the plasma membrane. Cells were then fixed with 3% paraformaldehyde in PBS and stained with affinity-purified rabbit anti-VI antibodies (16), Alexa Fluor 680-conjugated anti-mouse and Alexa Fluor 488-conjugated anti-rabbit antibodies (Thermo Fisher Scientific) and DAPI as previously described (21). Imaging was performed with a Leica SP5 confocal microscope as previously described (21). More information can be found in SI.

### Thermal stability assays

Fluorescence emission spectra from virus samples mixed with YOYO-1 were obtained every 2 ºC along a temperature range from 20 to 70 ºC in a Horiba FluoroLog spectrophotometer. The dye was excited at 490 nm and maximal emission intensity was achieved at 509 nm. More information can be found in SI.

### Mechanical stability assays

For AFM imaging, virus samples were diluted in a solution of NiCl_2_ in HBS to obtain a final solution of 5 mM of NiCl_2_ and virus concentrations of 1.5–2×10^11^ viral particles per ml. A drop of 20 µL of virus solution was deposited on freshly cleaved mica and incubated for 30 min at 4ºC before washing with 5 mM NiCl_2_ in HBS. The AFM (Nanotec Electrónica S.L., Madrid, Spain) was operated in jumping mode in liquid with the modifications described in (46), using RC800PSA (Olympus, Tokyo, Japan) cantilevers with nominal spring constants of 0.05 N/m. Monitoring virus disassembly in real time was accomplished by repeatedly imaging each particle as described (23). More information can be found in SI.

### Cryo-EM analyses

Methods to obtain the maps used in this work have been described elsewhere (33). These maps have been deposited at the Electron Microscopy Databank with ID EM-4448 (Ad5-wt) and EM-4424 (Ad5-VII-). To analyze capsid differences between Ad5-wt and Ad5-VII-particles, difference maps were calculated after filtering both maps at the same resolution (5.5 Å) using Xmipp and normalizing with UCSF Chimera. To delineate the extra density within the hexon cavities in Ad5-VII-, density maps at 5.5 Å resolution for each hexon trimer in the AU were created from the high resolution model (PDB ID = 6B1T) and subtracted from the Ad5-VII-map. More information can be found in SI.

## Acknowledgements

This work was supported by grant BFU2016-74868-P, co-funded by the Spanish State Research Agency and the European Regional Development Fund to CSM; as well as by grants from the Spanish Ministry of Economy, Industry and Competitiveness BIO2015-68990-REDT (the Spanish Adenovirus Network, AdenoNet) to CSM; FIS2017-89549-R, “María de Maeztu” Program for Units of Excellence in R&D (MDM-2014-0377), and FIS2017-90701-REDT to PJP. PH was funded by National Institutes of Health grants CA122677 and AI102577. UFG and MS were funded by the Swiss National Science Foundation (SNSF 31003A_179256/1) and the SNSF program Sinergia (CRSII5_170929/1). MM was funded by grant BFU2015-70052R, (MINECO/FEDER, UE) and the Ciber of Respiratory Diseases (CIBERES), an initiative from the Spanish Institute of Health Carlos III (ISCIII). MH-P is a recipient of a Juan de la Cierva postdoctoral contract funded by the Spanish State Research Agency. MP-I holds a predoctoral contract from La Caixa Foundation. We acknowledge the CNB-CIB cryo-EM facility for data acquisition, Roberto Marabini for assistance with image processing software, Milagros Castellanos and Luis A. Campos for advice with fluorescence measurements, Melanie Grove and Nicole Meili for negative stain EM of the Ad5-VII-preparation used in the immunofluorescence experiments, and Mara I. Laguna for expert technical help.

## Author Contributions

Designed research: U.F.G., P.H., P.J.P., C.S.M.

Specimen production: M.S., P.O., M.P.-I., G.N.C., P.H.

Entry assays: M. S., U.F.G.

Cryo-EM: M.H.-P., M.P.-I., G.N.C., C.S.M.

Thermostability assays: M.H.-P., M.P.-I., M.M.

Mechanical stability assays: N.M.-G., P.J.P.

Wrote the paper: M.H.-P. C.S.M., with input from all the authors.

