## Supplementary Information for "Dynamic competition for hexon binding between core protein VII and lytic protein VI promotes adenovirus maturation and entry"

**This PDF file includes:**

Supplementary methods

Figures S1 to S8

SI References

### Supplementary Methods

#### *Virus purification*

The Ad5-wt virus used in cryo-EM, AFM and extrinsic fluorescence assays was the E1-E3 deleted, GFP expressing Ad5/attP vector which is structurally wildtype (1), propagated at 37ºC in HEK293 cells and harvested at 36 h post-infection. The parent virus Ad5-VII-loxP with loxP sites flanking the protein VII open reading frame was generated and propagated as described (2). Ad5-VII- particles were produced using the Cre66 cell line, a Cre recombinase expressing cell line derived from HEK-293 cells (a gift from Dr. Stefan Kochaneck, University Ulm, Germany; unpublished), maintained in Dulbecco's modified Eagle's medium supplemented with 10% HyClone Fetalclone III serum, penicillin, streptomycin, and 0.25 mg/ml of Geneticin (Life Technologies). Cre66 cells (6 x $10^8$ total cells) were infected with 200 CsCl-purified Ad5-VII-loxP virus particles/cell for 1 h at 37 ºC. Infected cells were harvested 3 days after infection when full cytopathic effect was evident. The cells were pelleted by centrifugation, medium removed, and cell pellets flash frozen in liquid nitrogen for storage. Infected cell pellets were resuspended in DMEM and lysed by three or four freeze-thaw cycles. Cell lysates were clarified to remove cellular debris by centrifugation at 4000 rpm and 4 °C for 30 min. The pellet containing cell debris was discarded. For both Ad5-wt and Ad5-VII-, particles were purified by centrifugation at 18 °C and 217,485g in a 1.25-1.40 g/ml CsCl step gradient (90 min), followed by a 1.31 g/ml isopycnic one (18 h). CsCl solutions were prepared in TD1X buffer (137 mM NaCl, 5.1 mM KCl, 0.7 mM $Na_2HPO_4.7H_2O$, 25 mM Tris-HCl, pH 7.4). Bands extracted from the gradients were desalted on a Bio-Rad 10 DC column, and stored in HBS buffer (20 mM HEPES pH 7.8, 150 mM NaCl) supplemented with 10% glycerol at -80 °C until further use.

Capsid protein concentration was quantified using the hexon fluorescence emission spectra obtained in a Hitachi Model F-2500 FL spectrophotometer. Sample volumes of 0.150 ml were examined in sealed quartz cuvettes. The sample was excited at 285 nm, and the emission was monitored from 310 to 375 nm using excitation and emission slit widths of 10 nm. The spectra were corrected by subtraction of the buffer spectrum. The maximum emission intensity for each spectrum was found at 333 nm and recorded. The concentration was determined from a calibration curve obtained with a sample of known concentration (3).

Ad5-VII- particles used in entry assays were produced using A549 cells expressing a Cre recombinase (2) isolated and labeled with atto 565 as previously described (4-6), except that unincorporated atto 565 dye was removed with Zeba spin desalting columns (40K MWCO, ThermoFisher Scientific). The virus was stored in a buffer containing 10 mM Tris-HCl pH 8.1, 150 mM NaCl, 1 mM $MgCl_2$ and 10% glycerol at -80 °C until further use.

#### *Endocytosis and protein VI exposure assays*

A549 cells were seeded on coverslips at 40000 cells per well in a 24-well plate two days before the experiment. Ad5-wt and Ad5-VII- particles labeled with atto 565 were added to cells (MOI Ad5-wt ~ 62000 virus particles and Ad5-VII- ~ 149000 virus particles per cell) at 4 °C for 60 min in RPMI 1640 medium (without $NaHCO_3$) supplemented with 0.2% bovine serum albumin, 20 mM HEPES, and penicillin-streptomycin (RPMI-BSA medium). Unbound virus particles were removed, cells were washed once with cold RPMI-BSA medium and incubated in a 37 °C water bath in warm RPMI-BSA medium for 0, 10 or 20 min. Afterwards, intact cells were incubated with mouse 9C12 anti-hexon antibody in the RPMI-BSA medium at 4 °C for 1h to tag particles at the plasma membrane (the 9C12 antibody, developed by Laurence Fayadat and Wieber Olijve, was obtained from Developmental Studies Hybridoma Bank developed under the auspices of the National Institute of Child Health and Human Development). Cells were then fixed with 3% paraformaldehyde in PBS and stained with affinity-purified rabbit anti-VI antibodies (7) and Alexa Fluor 680-conjugated anti-mouse and Alexa Fluor 488-conjugated anti-rabbit antibodies (Thermo Fisher Scientific) and DAPI as previously described (8). Imaging was performed with a Leica SP5 confocal microscope as previously described.(8) Maximum projections of confocal stacks were analyzed by a custom-programmed MATLAB (The Mathworks) routine to score the number of plasma membrane-associated virus particles (particles with both the atto 565 and Alexa Fluor 680 signals) and internalized particles (particles with only the atto 565 signal) per cell, as well as protein VI signal on internalized virus particles (the MATLAB routine will be made available upon request). The threshold value for antibody-positive particles was determined by combining a virus image with an antibody image obtained from noninfected cells and taking the highest virus-associated antibody signal as a cutoff value. GraphPad Prism (GraphPad Software, Inc. La Jolla) was used for creating the scatter plots. Representative images shown in figures are maximum projections of confocal stacks, and the images were processed with Fiji applying the same changes in brightness and contrast to all image groups in the series.

#### *Thermal stability assays*

Virus samples ($5x10^9$ virus particles in a final volume of 800 µl of buffer) were incubated overnight at 4 °C in 8 mM $Na_2HPO_4$, 2 mM $KH_2PO_4$, 150 mM NaCl, and 0.1 mM EDTA, pH 7.4. A volume of 8.8 µl of YOYO-1 diluted 1:100 dilution in milliQ water from initial stock (Thermo Fisher Scientific, stock composition: 1 mM YOYO-1 in DMSO) was added to the virus sample and equilibrated for 5 min at 20 ºC before data acquisition. Fluorescence emission spectra were obtained employing a Horiba FluoroLog spectrophotometer equipped with a Peltier temperature control device. The dye was excited at 490 nm wavelength, and fluorescence emission was monitored from 500 to 580 nm using an excitation slit of 2 nm and an emission slit of 5 nm. Maximal emission intensity occurred at 509 nm. The temperature in the sample cell was increased from 20 to 70 ºC, with readings every 2 ºC after holding for 0.5 min for equilibration. Each run took approximately one hour and a half to be completed. Raw spectra were corrected by subtraction of the YOYO-1 spectrum in buffer at each tested temperature. The fractional change in fluorescence $F_n\ (T)$, was calculated according to equation (1):

$$F_n\ (T) = \frac{F_{exp} - F_i(T)}{F_f(T) - F_i\ (T)} \qquad (1)$$

Where T indicates the independent variable temperature (ºC) and $F_{exp}$ refers to the raw fluorescence signal (c.p.s) at each temperature. $F_i$ and $F_f$ represent the linear extrapolates, at this temperature, of pre-transition and post-transition base lines, respectively. A total of four independent experiments were considered for the averaged curve of each specimen. The half transition temperatures ($T_{0.5}$) were calculated from the fitting of the averaged fluorescence intensity curve as a function of temperature (T) to a Boltzmann sigmoid equation, $F_s(T)$ (Origin software package) according to equation (2):

$$F_S(T) = F_{max} + \frac{F_0 - F_{max}}{1 + \exp\left(\frac{T - T_{0.5}}{dT}\right)} \qquad (2)$$

***Mechanical stability assays***

For AFM imaging, virus samples were diluted in a solution of $NiCl_2$ in HBS to obtain a final solution of 5 mM of $NiCl_2$ and virus concentrations of 1.5–2x$10^{11}$ viral particles per ml. A drop of 20 µL of virus solution was deposited on freshly cleaved mica and incubated for 30 min at 4 ºC before washing with 5 mM $NiCl_2$ in HBS. The AFM tip was pre-wetted, and the mica was placed on a holder and immersed in 500 ml of the same buffer. The virus is therefore maintained in a hydrated, close to physiological state throughout the experiment. The AFM (Nanotec Electrónica S.L., Madrid, Spain) was operated in jumping mode in liquid with the modifications described in (9), using RC800PSA (Olympus, Tokyo, Japan) cantilevers with nominal spring constants of 0.05 N/m. Cantilevers were routinely calibrated using Sader's method (10). Images of 128x 128 points and 300 x 300 $nm^2$ were recorded scanning from left to right by applying forces of 100 pN at a temperature of 20 ºC to avoid thermal drift. Monitoring virus disassembly in real time was accomplished by repeatedly imaging each particle as described (11). To analyze volume changes, all images corresponding to the same virion were aligned and flooded using the WSXM software (http://www.wsxm.es/). The flooding height was determined for extracting only the volume contribution of the viral particle or major capsid debris, not of single capsomers spread on the substrate once disassembly started. Volume measurements were normalized to the initial value for each individual particle. The volume loss rate (given in volume percentage per image) was estimated from the slope of the volume loss curves after the three pentons in the upper particle facet had been released.

***Cryo-EM analyses***

Methods to obtain the maps used in this work have been described elsewhere (12). Briefly, purified Ad5-wt and Ad5-VII- particles were vitrified in glow discharged Quantifoil R2/4 300 mesh Cu/Rh grids using a Leica CPC plunger. Cryo-EM images (4395 for Ad5-wt, 1640 for Ad5-VII-) were recorded at the CNB-CSIC cryoEM facility in a 200 kV Talos Arctica microscope (FEI) equipped with a Falcon II detector, with a total dose of 50 e-/$Å^2$ distributed over 32 frames, at a nominal pixel size of 1.4 Å and defocus range between -0.5 and -3 µm. All image processing and 3D reconstruction tasks were performed within the Scipion framework (13). Frames were aligned using Motioncor2 and weighted according to the electron dose received before averaging (14). The CTF was estimated using CTFFIND4 (15). Micrographs were corrected for the phase oscillations of the CTF (phase-flipped). Particles (26661 for Ad5-wt, 27909 for Ad5-VII-) were extracted into 752×752 pixel boxes, normalized and resized to a sampling rate of 2.8 Å/px (376 px box size), using Xmipp (16). All 2D and 3D classifications and refinements were performed using RELION (Scheres 2012). 2D and 3D classification were used to discard low quality particles. Icosahedral symmetry was imposed throughout 3D classification and refinement. The class yielding the best resolution map (23060 particles for Ad5-wt, 21762 particles for Ad5-VII-) was individually refined using the original 752 px boxed particles, yielding a final map at 5.5 Å resolution for Ad5-wt and 5.0 Å for Ad5-VII-. Resolution was estimated according to the gold-standard FSC = 0.143 criterion as implemented in RELION auto-refine and postprocess routines (17). These maps have been deposited at the Electron Microscopy Databank with ID EM-4448 (Ad5-wt) and EM-4424 (Ad5-VII-). High frequencies were boosted by using a temperature factor of -363 $Å^2$ (Ad5-wt) or -440 $Å^2$ (Ad5-VII-). The actual sampling for both maps was estimated using UCSF Chimera *Fit in Map* (18), by comparison with the HAdV-C5 high resolution model (PDB ID = 6B1T) (19), yielding a value of 1.37 Å/px.

To analyze the capsid differences between Ad5-wt and Ad5-VII- particles, difference maps were calculated after filtering both maps at the same resolution (5.5 Å) using Xmipp and normalizing with UCSF Chimera (*vop scale # sd 1*). To delineate the extra density within the hexon cavities in Ad5-VII-, density maps at 5.5 Å resolution for each hexon trimer in the AU were created from the high resolution model (PDB ID = 6B1T) using Chimera *molmap*, and subtracted from the Ad5-VII- map (*vop subtract min RMS*). α-helices of different lengths were modeled with Coot (20).

### Supplementary figures

**Figure S1: Cartoon illustrating the relative sizes of the adenovirus maturation protease (AVP) and the hexon cavity, as well as the location of the pVI$_N$ cleavage site.** Hexon 2 is shown in pink ribbon. The pVI$_N$ peptide is colored cyan and the central fragment of pVI is in green (19). AVP (PDB ID: 1AVP) is in blue (21).

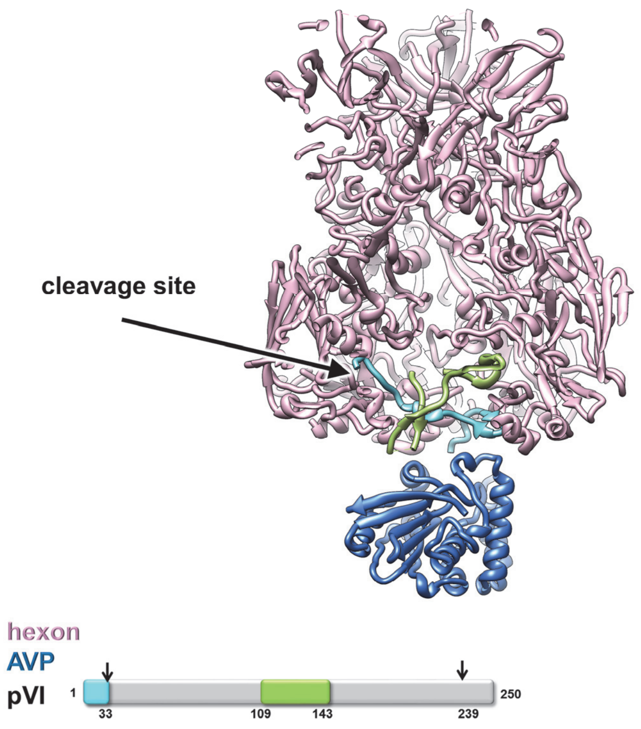

**Figure S2: Negative staining electron microscopy of the Ad5-VII- preparations used for the immunofluorescence experiments (a) or for cryo-EM, AFM and extrinsic fluorescence assays (b)**. Some disrupted particles are present in (a), consistent with the observation of signal for protein VI before allowing cell entry (related to Fig. 1b)

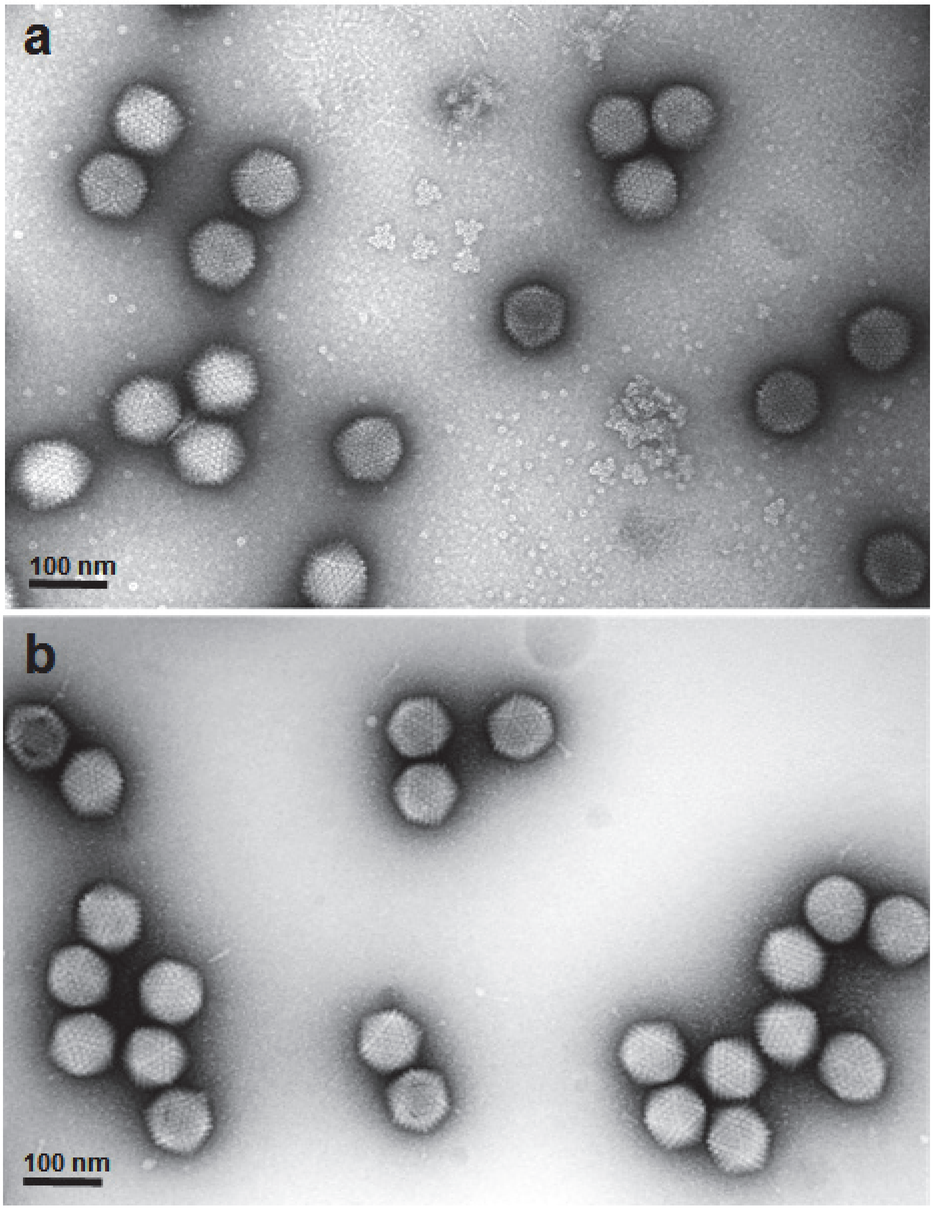

**Figure S3**: **Both Ad5-wt and Ad5-VII- particles are efficiently endocytosed.** The dataset is the same as in Fig.1. The scatter plot shows the fraction of 9C12 antibody-negative virus particles per cell at the indicated time points, one dot representing one cell. Error bars represent the means ± SD.

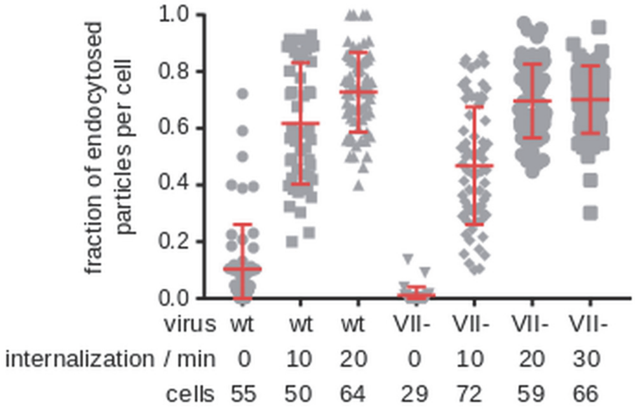

**Figure S4: Ad5-wt and Ad5-VII- cryo-EM maps.** Left column: representative cryo-EM micrographs of Ad5-wt and Ad5-VII-. Right column: detail of the central sections of the cryo EM density maps of Ad5-wt and Ad5-VII-. Higher density is shown in white. A white line indicates the position of a 5-fold icosahedral symmetry axis in Ad5-wt. The white circle indicates extra density within the hexon cavity in the Ad5-VII- map.

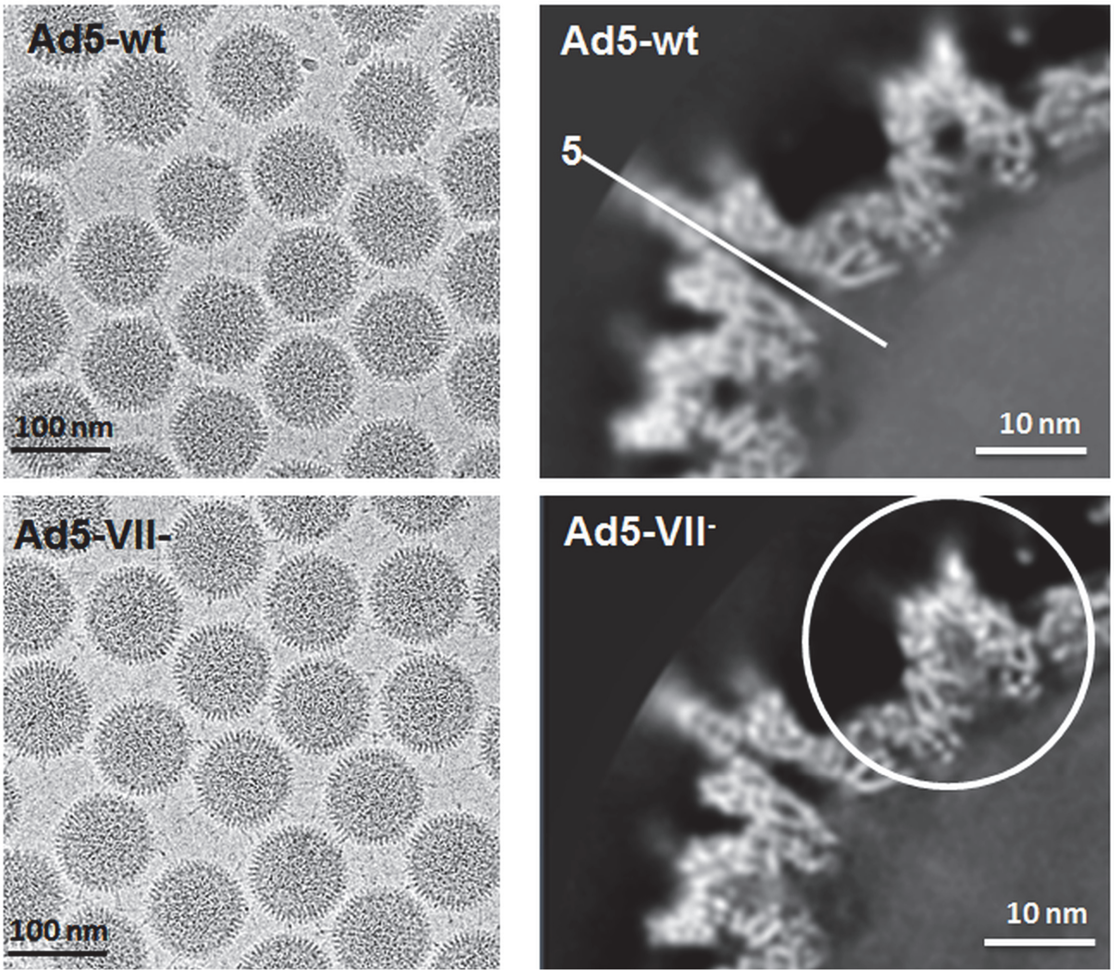

**Figure S5**: **Cryo-EM difference map showing density present in Ad5-wt but absent in Ad5-VII- particles, contoured at several different thresholds.** The four hexon trimers in the AU (see inset) as seen from inside the viral particle are shown as a gray surface generated from the high resolution structure (PDB 6B1T). In the two bottom panels, the difference map obtained by subtracting the two cryo-EM maps (Ad5-wt minus Ad5-VII-) is shown as a semitransparent red surface contoured at different thresholds, as indicated. Red dashed circles indicate difference densities located at positions in the hexon cavity where either $pVI_N$ or $pVII_{N2}$ would bind, according to Dai et al. Cyan, green and orange ribbons correspond to the pVI and pVII fragments traced by (19), as indicated in the schematics in Fig. 4c. The inset shows a schematic of the icosahedral asymmetric unit as seen from inside the particle, with the four hexon trimers numbered 1-4, and the positions of the 2, 3 and 5-fold icosahedral symmetry axes depicted by a red oval, triangle and pentagon, respectively.

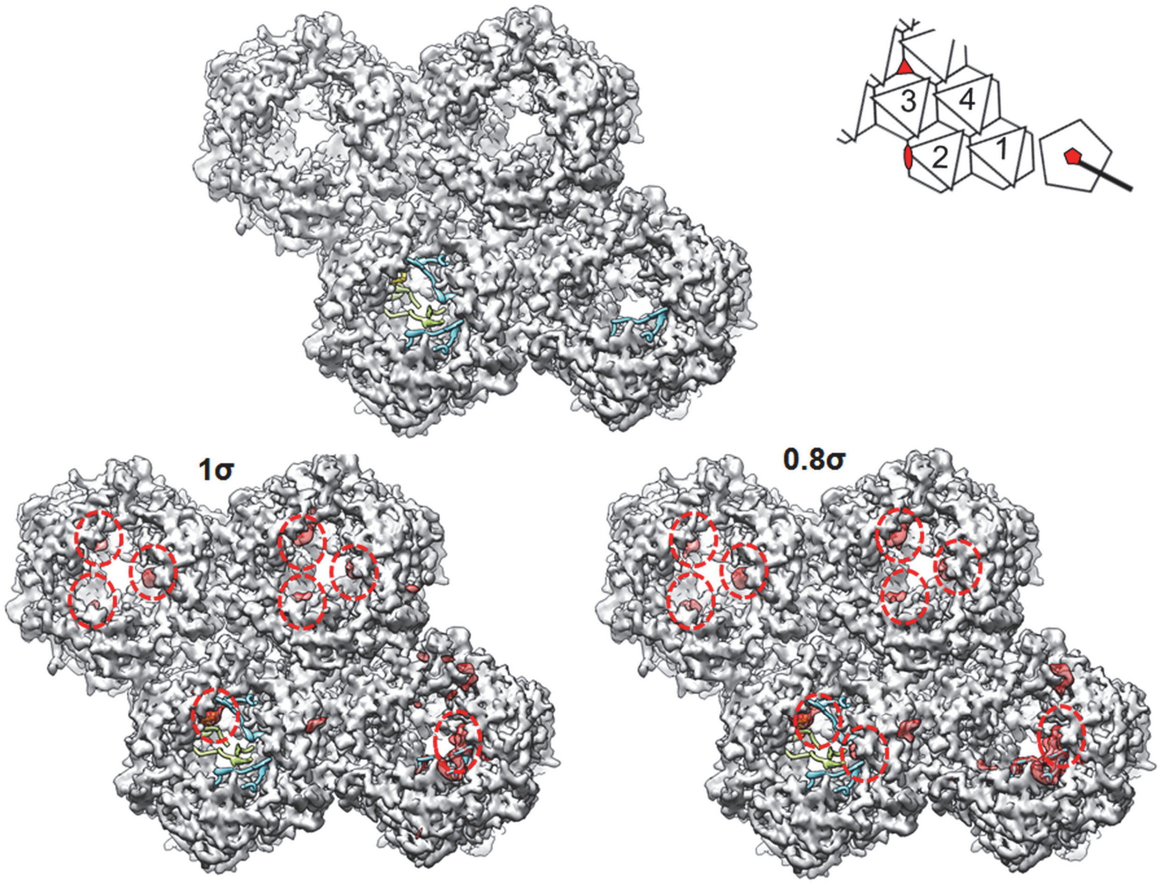

**Figure S6: Cryo-EM difference map showing density present in Ad5-VII- but absent in Ad5-wt particles, contoured at several different thresholds.** The four hexon trimers in the AU (see inset) as seen from inside the viral particle are shown as a gray surface generated from the high resolution structure (PDB 6B1T). The difference map obtained by subtracting the two cryo-EM maps (Ad5-VII- minus Ad5-wt) is shown as a blue surface contoured at different thresholds, as indicated. Cyan, green and orange ribbons correspond to the pVI and pVII fragments traced by (19), as indicated in the schematics in Fig. 4c. The inset shows a schematic of the icosahedral asymmetric unit as seen from inside the particle, with the four hexon trimers numbered 1-4, and the positions of the 2, 3 and 5-fold icosahedral symmetry axes depicted by a red oval, triangle and pentagon, respectively.

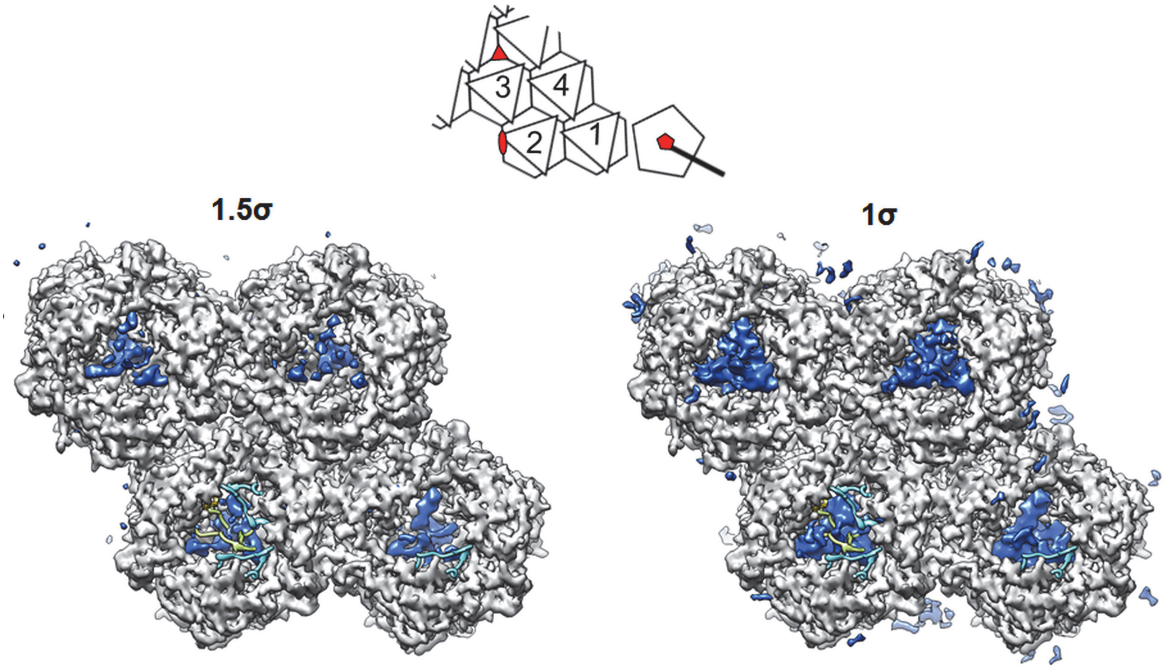

**Figure S7: Secondary structure prediction for pVI.** The secondary structure prediction for pVI was obtained using Jpred (22). Only the N-terminal part of the protein (residues 1-120) is shown here. Red bars represent the predicted formation of α-helices. The position of the predicted lytic peptide (23) is indicated (arrows).

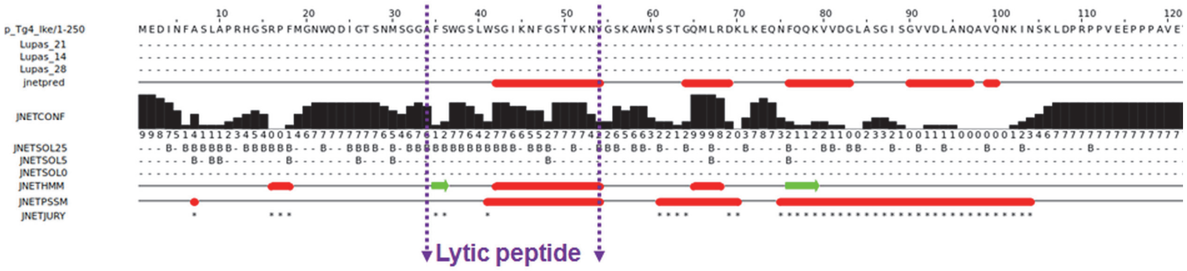

**Figure S8: Density corresponding to partially uncleaved pVI inside the cavity of all hexon trimers in the asymmetric unit.** The four hexon trimers in the AU (H1-H4) are represented as pink ribbons. In gray mesh, the difference density obtained by subtracting the density for each hexon trimer, calculated at 5.5 Å resolution from PDB 6B1T, from the Ad5-VII- cryo-EM map. The difference maps are contoured at 1.7σ. Three copies of the $pVI_N$ peptide (one in H1 and two in H2) as traced in PDB 6B1T are represented as cyan ribbons. Panels at the right hand side of each hexon show only the difference density and the $pVI_N$ ribbons, for a clearer view. Only the region of the difference map contained within 12 Å from helices modeled in the tubular densities within the hexon cavity is shown.

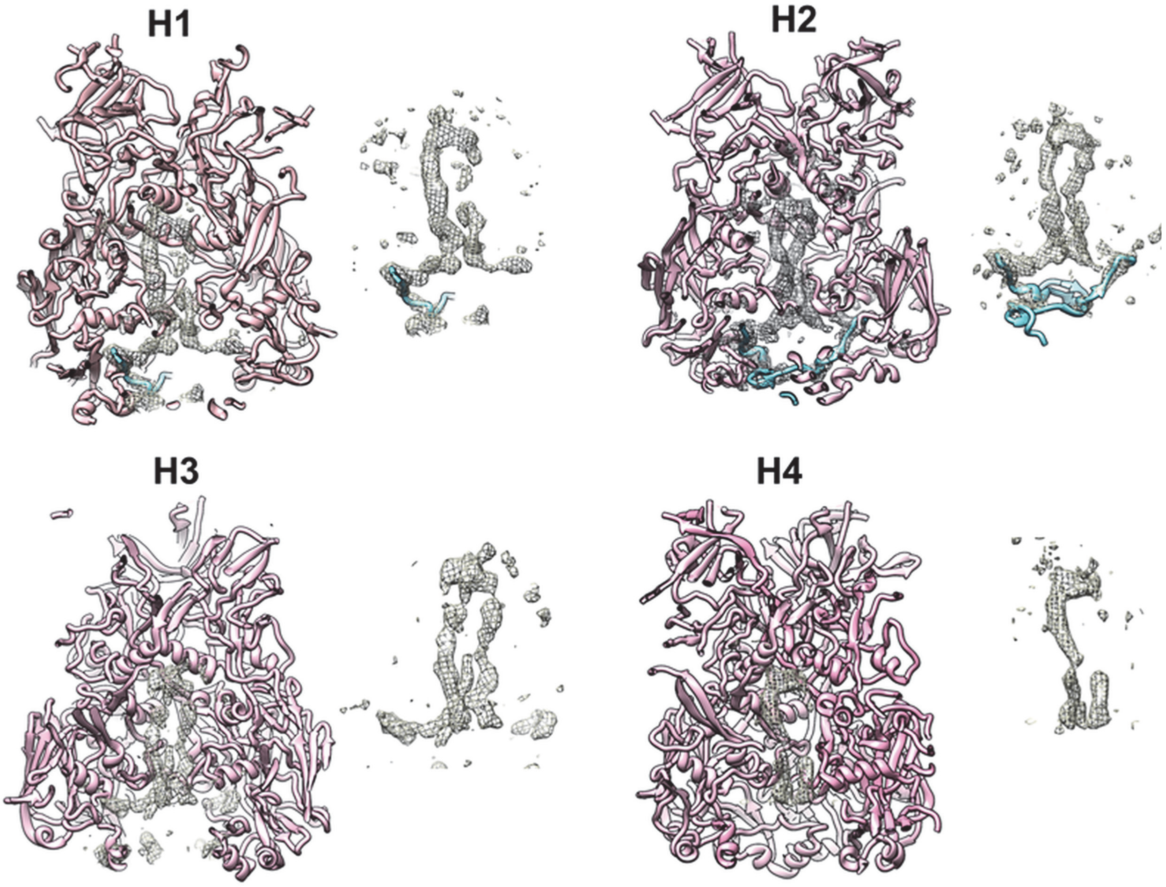
